# A Statistical Approach to Cellular Resource Allocation Models

**DOI:** 10.64898/2026.08.15.744912

**Authors:** Ankita Roychoudhury, David Pincus, Madhav Mani

## Abstract

Understanding how cells regulate growth despite molecular complexity remains a central question in quantitative biology. While thousands of genes respond to environmental perturbations, the population growth rate varies smoothly across conditions, suggesting the existence of simple organizing principles. Here, we show that statistical analysis of mRNA composition across environmental conditions reveals growth tradeoffs across organisms including *E. coli* and *S. pombe*. Using partial least squares regression, we identify two opposing gene sectors whose coordinated expression encodes growth rate. A minimal transcription-translation model, constrained by empirical scaling laws of total mRNA and ribosomal fractions, explains this tradeoff as a necessary consequence of the empirical observations. Extending the model to include charged tRNA dynamics reveals distinct regulatory regimes: *E. coli* operates co-limited by ribosomal mRNA and charged tRNA availability, whereas *S. cerevisiae* is primarily ribosomal mRNA-limited. Together, these results provide a statistical method to determine key tradeoffs across organisms and offer a framework to interpret organism-specific growth regimes.

## I. INTRODUCTION

The process of cellular homeostasis involves the adaptation of a cell’s internal processes in response to external stimuli. Upon recovering homeostasis in an altered environment, cells adopt reproducible growth rates that are continuous functions of environmental parameters (Fig. 1A). A fundamental challenge in biology is understanding how highly diverse cell types, despite vast differences in physiology and environment, regulate growth through shared intracellular principles (Fig. 1B).

**FIG. 1.**
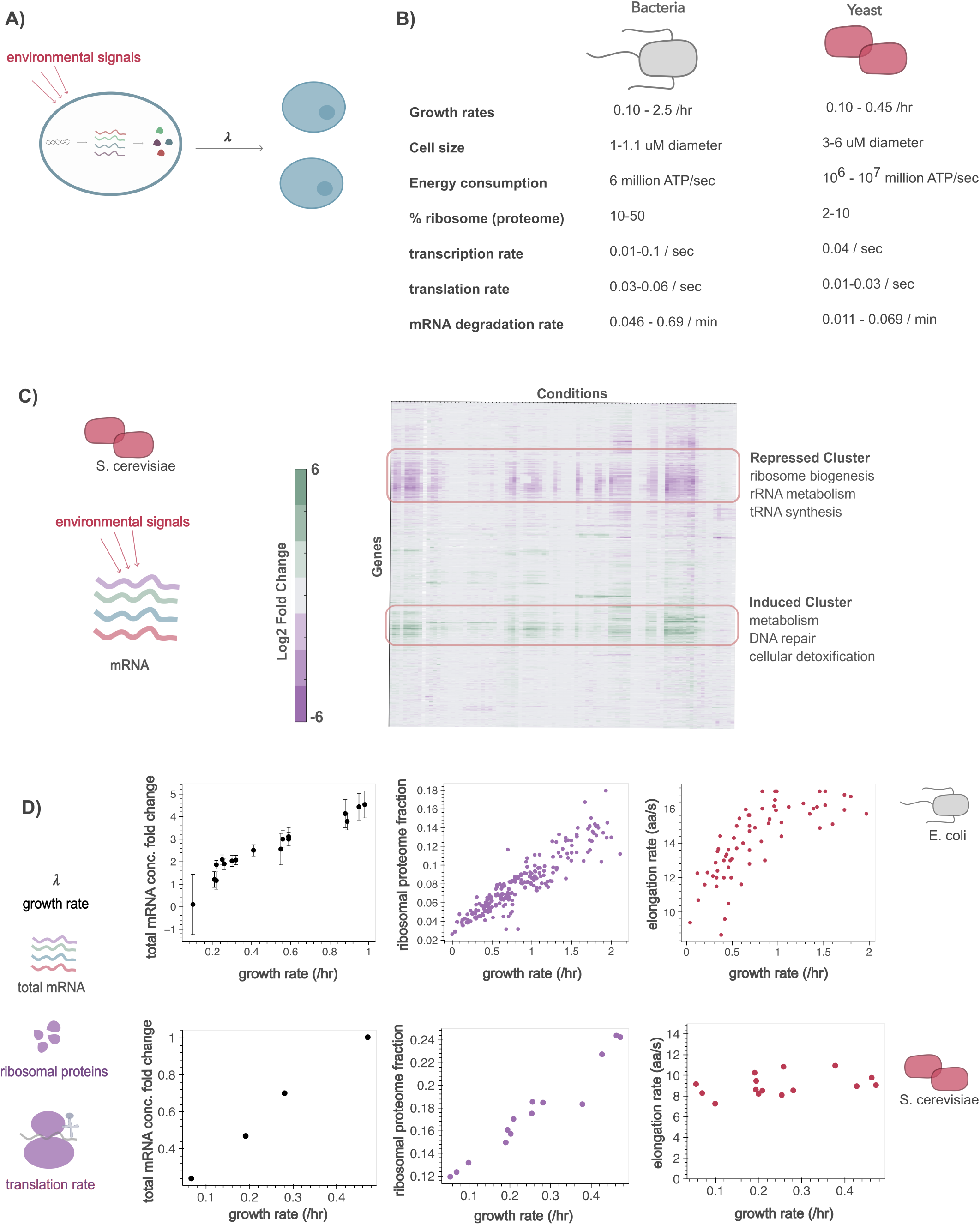
Existing theories and data around growth regulation. A) Cells adapt to environmental signals via the central dogma. This, in turn, affects the steady state growth rate of the cell. B) Key biophysical parameters for bacteria and yeast. Parameter values are from references [2, 3, 17–21]. C) Existing data exploring mRNA composition. Heatmap adapted from Gasch et al. [8]. The gene expression profile that defines the Environmental Stress Response (ESR). D) Existing theories and data around total mRNA, ribosome proteome fractions, and elongation rates across growth rates. The top data panels represent work in *E. coli*. (Balakrishnan et al. and Chure et al.) The bottom data panels are adapted from *S. cerevisiae* (Gao et al.) using automeris.io

A purely molecular perspective of a cell sees it as a collection of thousands of genes, proteins, and metabolites, along with tightly regulated rates of transcription, translation, and post-translational modifications. This emphasizes how specific environmental inputs cause specific molecular responses within the cell. Despite this manifest molecular complexity, multiple studies, mostly in model prokaryotic and single-cell eukaryotic systems, have revealed simpler dynamics in cellular responses across diverse perturbations [1–4]. In *E. coli*, quantitative proteome measurements, coupled with coarse-grained grouping of proteins into sectors, provide a predictive physiological rationale for the variation in growth rates across environments [1,5–7]. This framework rests on a simple constraint: because the proteome represents a finite resource, growth-related genes partition into sectors — most notably ribosomal and metabolic — whose allocations must trade off against one another. This constraint alone, independent of the regulatory logic of individual genes, is sufficient to predict quantitative relationships between sector composition and growth rate. Though far from a complete picture, these studies put forward a complementary integrated framework, describing growth regulation in terms of functionally defined sectors of a cell’s processes rather than the specific logic of individual pathways.

Although substantial work has clarified the relationship between protein composition and growth rate, comparable studies of mRNA have largely focused on total transcript abundance [2,3] or qualitative shifts in expression under stress [8], leaving open how the relative **composition** of the transcriptome quantitatively relates to growth rate across conditions [7]. This is particularly relevant in cells whose growth rate responds to regulatory signals beyond nutrient limitation cues. A seminal study conducted by Gasch et al. subjected *S. cerevisiae* to various stressors and measured mRNA abundance across cell populations [8]. The conditions spanned a wide variety – including heat shock, amino acid starvation, nitrogen depletion, hydrogen peroxide treatment, hyper-osmotic shock, and more – allowing for the exploration of a general transcriptional response across conditions. Through statistical analyses of their data, they identified that ~ 15% of genes are consistently induced or repressed 16-fold relative to an unstressed condition across multiple stress conditions (Fig. 1C) [8]. Upon analyses of these groups, the genes that are repressed are related to ribosome biogenesis, rRNA metabolism, and tRNA synthesis while the induced genes are related to metabolism, DNA repair, and cellular detoxification. While a direct link has not yet been established between these stress-induced gene expression responses and the growth rate, this study elucidates the potential low-dimensionality of the transcriptional response to perturbation. Despite the diversity of stressors, a consistent core of genes shifts together, hinting at an intrinsic, growth-coupled program.

Increasing evidence indicates that total mRNA concentration is also closely coupled to growth. Studies in *E. coli* and *S. cerevisiae* reveal positive linear relationships between total mRNA concentration and growth [2,3]. Balakrishnan et al. measured total mRNA concentrations in different conditions, including carbon limitation, ammonia limitation, and translation inhibition [3]. They find a linear relationship between total mRNA concentration and growth (Fig. 1D). A similar relationship was reported by Gao et al. in *S. cerevisiae* across four distinct carbon sources [2]. In addition, *E. coli* and *S. cerevisiae* exhibit a positive relationship between the ribosomal fraction of the proteome and growth rate in both works (Fig. 1D). These observations highlight a consistent pattern: faster-growing cells contain proportionately more ribosomes and higher total mRNA levels. The emergence of these simple linear relationships between cellular composition and growth rate, despite the enormous complexity of the cell, is what motivates the community search for simple organizing principles.

To understand these simple empirical relations, theoretical and experimental studies have sought to identify the fundamental limiting factor that sets growth rate across organisms. Classic bacterial growth models posit that growth rates are governed not just by ribosome abundance but also by the dynamics of translation, including the supply of charged tRNAs that deliver amino acids to elongating ribosomes [5,6,9]. In *E. coli*, this coupling means peptide elongation rate itself scales with growth rate, providing a natural explanation for how the quality and availability of environmentally sourced nutrients set the rate of proliferation. In contrast, recent work challenges this picture in eukaryotes: Gao et al. find that, like *E. coli*, ribosome concentration scales linearly with growth rate in *S. cerevisiae*, but unlike *E. coli*, peptide elongation speed remains constant at ~ 9 amino acids per second (Fig. 1D) [2]. Because elongation rate is itself set by the concentration of charged tRNAs, this constancy indicates that charged tRNA supply is not limiting translational capacity in *S. cerevisiae*. Instead, since ribosome production scales with growth without a corresponding change in elongation kinetics, the limiting step must lie upstream of elongation. They therefore propose that growth control is more likely governed by the formation of productive mRNA–ribosome complexes, rather than by deficits in charged tRNAs. Consistent with this view, theoretical work by Calabrese et al. proposes a quantitative framework in which growth is determined by ribosome–mRNA complex formation and can be limited by either total mRNA abundance or ribosome availability depending on cellular state [10]. In the intermediate regime, growth emerges from a trade-off between mRNA supply and ribosome allocation, while in limiting regimes growth becomes dominated by either transcriptional output or translational capacity. Thus, whereas prokaryotic growth models often emphasize translation-elongation dynamics and tRNA charging as central constraints on growth, eukaryotic systems appear to operate under a different regime, where mRNA availability and its interaction with ribosomes play a more dominant role.

If mRNA availability is indeed the limiting factor on translation in eukaryotes, then the composition of the transcriptome — which genes are expressed, and in what relative proportions — becomes a central determinant of growth itself. The established relationships between total mRNA abundance, ribosome content, and growth rate motivate a deeper examination of the transcriptome. What is the origin of the simple compositional structure in the yeast transcriptome across stress conditions and what is its relationship with growth rate? Here, we ask whether statistical analysis of mRNA composition can identify the resource-allocation tradeoffs that govern growth, and whether a minimal biophysical model can explain why these tradeoffs must exist. Such a framework offers a data-driven way to identify regulatory sectors without presupposing gene function in advance. It also clarifies which resource limits growth in a given organism, explaining why growth strategies diverge across species and enabling predictions of how growth responds to perturbation. Using datasets from prokaryotes (*E. coli*) and eukaryotes (*S. pombe* and *S. cerevisiae*) [3,11], we first show that mRNA composition encodes growth rate through organism-specific tradeoffs. We then demonstrate that empirical scaling laws impose constraints that necessitate this tradeoff. Finally, we extend the model to include charged tRNA dynamics, revealing that distinct growth-limiting regimes emerge naturally from the same underlying framework.

## II. RESULTS

### A. mRNA composition scales with growth in *S. pombe* and *E. coli*

Kleijn et al. grew *S. pombe* in turbidostats with eight distinct nitrogen sources in triplicate [11]. Nitrogen was supplied either as ammonium chloride or as a single amino acid replacing ammonium as the primary nitrogen source. They measured growth rates, mRNA levels with mRNA-sequencing, and protein levels with mass-spectrometry (Fig. 2A), allowing us to investigate the relationship between mRNA composition and growth. Variation in the nitrogen-rich amino acids produces a growth rate range from 0.05 to 0.28 hr^−1^. Overall, we observe a linear trend between mRNA counts and protein abundances in log-log space (r = 0.61), consistent with prior reports of moderate mRNA–protein correlation in yeast [12,13] (Fig. 2A). Mirroring Gasch et al. we performed hierarchical clustering on the mRNA-seq data from Kleijn et al., which provides insights into the large-scale patterns in the transcriptomic response [8]. This analysis takes the ammonium chloride conditions with the highest growth rate as the reference condition. Figure 2B highlights that 18% of genes are induced and 21% of genes are repressed independent of the nitrogen source. Genes repressed under stress are enriched for ribosome biogenesis and translation, while genes induced include meiosis and amino acid transport. In a manner similar to what Gasch et al. observed in *S. cerevisiae* (Fig. 1C), the results in *S. pombe* indicate a low dimensional structure in the transcriptomic response to environmental perturbations that is intrinsic to *S. pombe* and not highly dependent on the types of environmental perturbations.

**FIG. 2.**
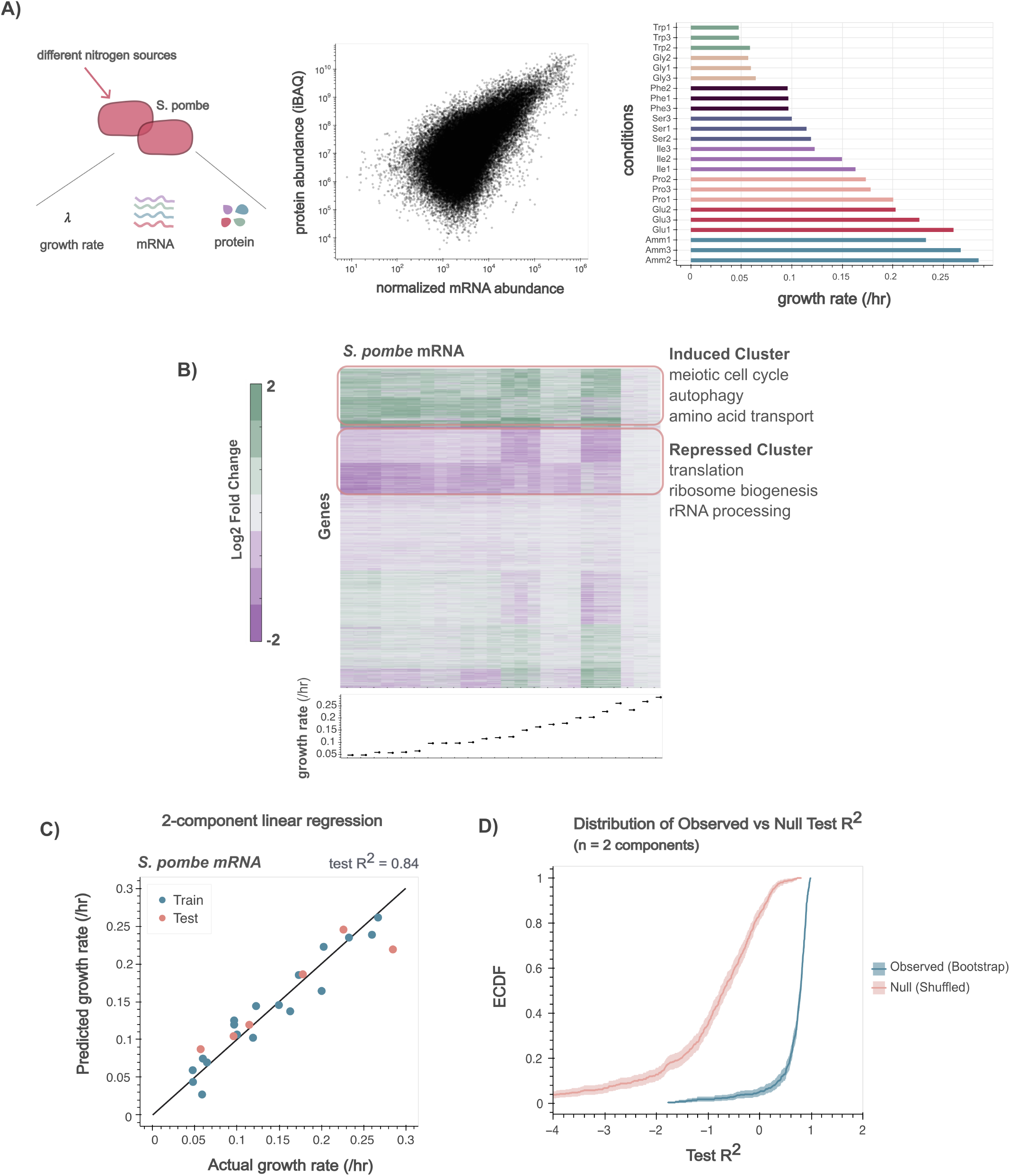
Data from *S. pombe* (Kleijn et al.) shows the relationship between mRNA composition and growth. A) Kleijn et al. collected data in *S. pombe* grown in media with 8 different nitrogen sources and 3 replicates. Proteomics, mRNA sequencing, and growth data was collected. Left plot shows protein abundance vs mRNA counts has the expected predictability. The right plot shows the range of growth rates captured, within replicates and across conditions. B) A hierarchical clustering of the mRNA-sequencing data shows the existence of genes that are induced and repressed independent of the nitrogen source used. C) PLS on the mRNA-sequencing data shows that growth information is indeed encoded in the transcriptome. D) Model validation via permutation testing. ECDF of test *R*^2^ values for the n = 2 component model across 500 bootstrap replicates. The observed bootstrap distribution is significantly shifted from the null distribution of shuffled labels, confirming the model captures genuine biological signal. 95 % confidence intervals are shaded. X-axis is truncated to [−4, 2] for visual clarity.

It is valuable to juxtapose the analysis in *S. pombe* to *E. coli*, to reveal the qualitatively distinct nature of growth regulation. In particular, since *E. coli* has been shown to be environmentally limited through its determination of the limiting pool of charged tRNAs, we anticipate the lack of a coherent grouping of induced and repressed genes across all conditions. Instead, we anticipate strong dependencies of the transcriptomic response on the nature of the environment [14]. We reanalyze data collected by Balakrishnan et al. in *E. coli* where they measured growth rates, mRNA levels via bulk mRNA-sequencing, and protein levels via mass spectrometry in carbon-limited, ammonia-limited, and translation-inhibited conditions (Fig. S1A) [3]. Hierarchical clustering of the *E. coli* data reveals a lack of a general transcriptomic pattern (Fig. S1B). Instead of a global transcriptomic pattern independent of the condition, strong vertical striations indicate condition-specific responses. This suggests that *E. coli* responds to perturbations via extrinsic regulation. This contrasts with the intrinsic response observed in *S. pombe* and *S. cerevisiae*, where transcriptomic variation is dominated by a conserved program across environmental conditions.

To quantify the relationship between mRNA composition and growth, we performed partial least squares (PLS) regression to identify linear combinations of genes that are predictive of growth rate (Fig. 2C). PLS models growth as a weighted sum of standardized gene expression values,

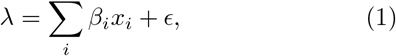

where *x*_*i*_ is the standardized expression of gene *i, β*_*i*_ is its PLS-derived coefficient, and *ϵ* is residual error. PLS extends PCA by identifying directions of covariation that are maximally predictive of growth rate, rather than directions of maximum variance alone. In *S. pombe*, the PLS model trained on 75% of the data predicted growth in the remaining 25% with *R*^2^ = 0.84. We determined the optimal number of PLS components (*n* = 2) by ensuring that additional components do not lead to overfitting (Fig. S2).

To guarantee robustness of the model, we compared its performance against a null distribution generated by shuffling growth rate labels across 500 permutations. Across multiple random seeds and cross-validation splits, the model trained on biological data yielded substantially lower errors and higher *R*^2^ values than the null models (Fig. 2D, S2A). Together, these results demonstrate transcript abundances are coordinated and predictive of growth in *S. pombe* without considering protein levels or post-translational regulation. This raises the question: What does the low-dimensional structure revealed by our statistical analyses tell us about resource allocation?

### B. What does PLS reveal about resource allocation models?

As introduced above, PLS regression yields coefficients (*β* values, Eq. 1) quantifying each gene’s contribution to predicting growth rate. Each PLS mode allows us to identify genes with positive and negative coefficients that covary with one another in a manner predictive of growth. Prior work identified a total proteome constraint in *E. coli*, where cells inhabit distinct growth states along a Pareto front defined by ribosomal and metabolic functional sectors [11]. Motivated by this, we ask whether our statistical model can reveal analogous cellular tradeoffs in a data-driven manner across novel cellular contexts (Fig. 3A).

**FIG. 3.**
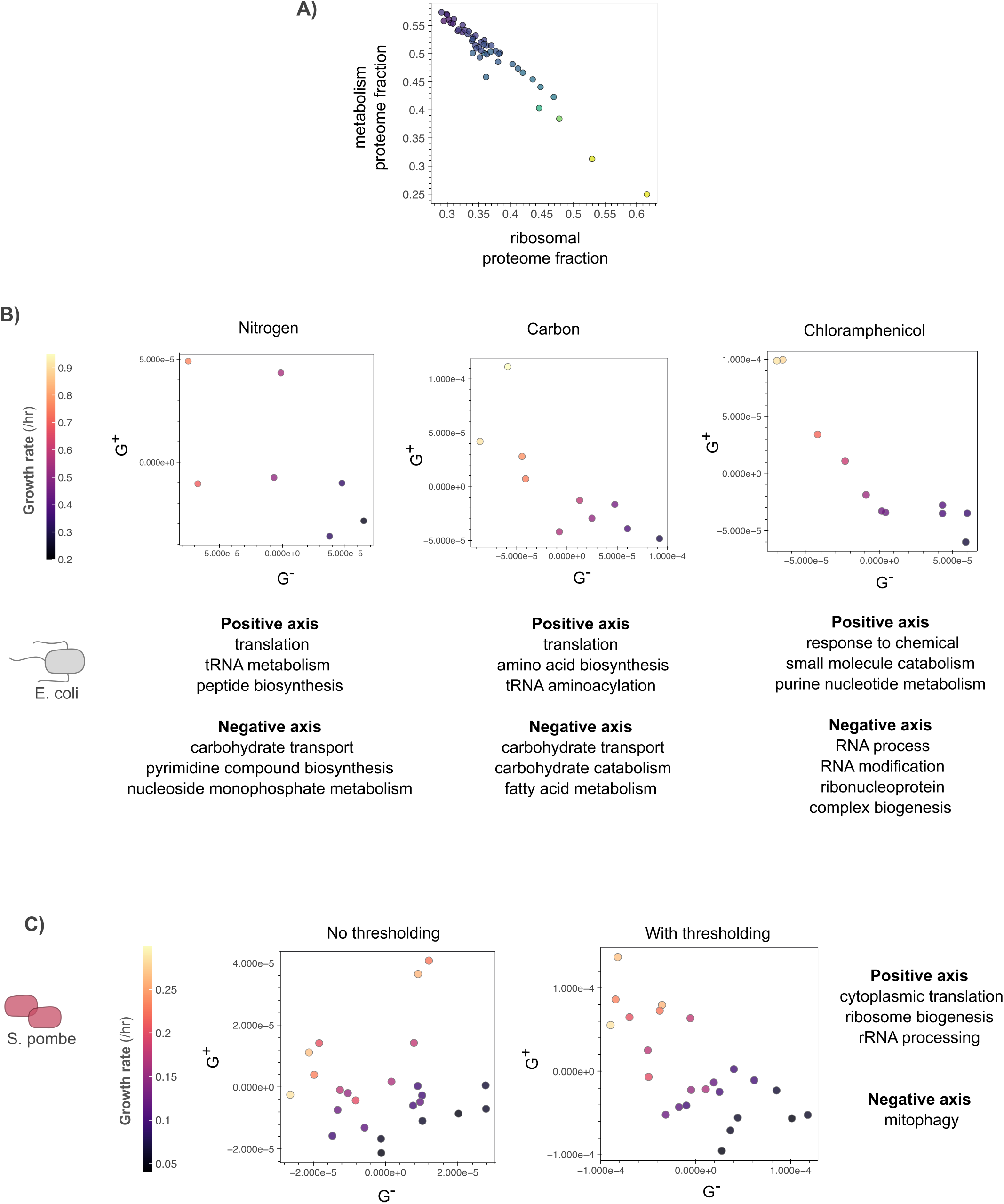
Tradeoff axes defined by PLS regression coefficients. A) Theoretical resource allocation model from Hwa et al. shows the tradeoff between ribosomal and metabolic proteome fraction across growth rates in *E. coli*. Data from [22] and plot adapted from [23]. B) PLS-derived tradeoff axes for *E. coli* across three nutrient limitation conditions. C) PLS-derived tradeoff axes for *S. pombe* across different nitrogen sources. Each point represents a sample, positioned by its projection onto the positive (growth-promoting genes, y-axis) and negative axis (growth-suppressing genes, x-axis), and colored by growth rate. GSEA reveals annotations for the positive and negative axis.

The distribution of *β* coefficients differs markedly between organisms (Fig. S3A). In *S. pombe*, where PLS was fit across all conditions simultaneously, a small number of genes exhibited large *β* values – consistent with a conserved stress response where the same genes predict growth across diverse perturbations. Permutation testing (500 shuffles of growth labels) identified a statistical threshold (*τ* = 6 *×* 10^−5^) above which *β* coefficients were significant (FDR *<* 0.05). This threshold identified 22 growth-promoting genes, G^+^ (*β > τ*), and 4 growth-suppressing genes, G^−^ (*β <* − *τ*). In contrast, *E. coli β* coefficients within each condition showed broader distributions (Fig. S3A), with fewer genes reaching extreme values. This may reflect the smaller sample size per condition (N=10) or a genuinely more distributed regulatory architecture. Rather than applying a statistical threshold for *E. coli*, we used the sign of *β* within each condition to partition genes: those with *β >* 0 define the growthpromoting axis (G^+^), while those with *β <* 0 define the growth-suppressing axis (G^−^).

For each sample, projection scores onto the G^+^ and G^−^ axes were computed as weighted sums of standardized gene expression values, where the weights are the PLS *β* coefficients restricted to genes in G^+^ or G^−^ respectively:

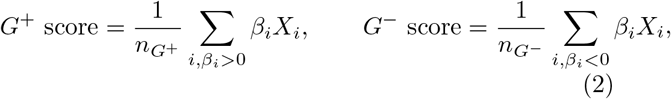

where *X*_*i*_ is the standardized expression of gene *i*, and 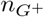 and 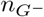 are the number of genes in each group. Plotting the G^+^ score against the G^−^ score for each sample thus positions each sample in a 2D space defined by the growth-promoting and growth-suppressing transcriptional programs (Fig. 3B, C). In *S. pombe*, the projection uses only the 26 statistically significant genes; in *E. coli*, all genes are used. This approach reveals the tradeoff geometry in a data-driven manner, without assuming the biological identity of gene sets *a priori*. A natural critique of this approach is the linearity of the statistical model that maps gene expression to growth rates, and thus our results could be seen as a first-pass with limited data. Advanced nonlinear approaches could be applied to this problem given more data.

To interpret the biological meaning of the PLS-derived axes, we performed gene set enrichment analysis (GSEA) on G^+^ and G^−^ (Fig. S3). GSEA takes genes ranked by *β* coefficients and tests whether predefined functional gene sets are enriched among genes with large positive or negative *β* coefficients. In *E. coli*, GSEA revealed the positive axis (G^+^) is consistently enriched for core growth functions – including translation, amino acid biosynthesis, and tRNA metabolism – across both carbon and nitrogen limitation. In contrast, the negative axis (G^−^) captures condition-specific programs. Under nitrogen limitation, G^−^ is enriched for nucleotide biosynthesis whereas under carbon limitation, G^−^ is enriched for carbohydrate transport and catabolism (Fig. 3B). For chloramphenicol, a translation inhibitor, the positive axis includes response to chemical while the negative axis is enriched for RNA modification and ribosome biogenesis. Full GSEA results are shown in Figure S3. These findings indicate that PLS recovers the canonical ribosomal-metabolic tradeoffs under nutrient limitation, with the negative axis reflecting condition-specific adaptations. This protocol can then be used to explore the most salient cellular tradeoffs in other organisms.

Performing a similar analysis in the *S. pombe* dataset, we observed that the positive axis is enriched for cytoplasmic translation and ribosome biogenesis while the negative axis is enriched for mitophagy (Fig. 3C). Enrichment of the negative axis for mitophagy suggests that slower-growth states in *S. pombe* are associated with increased investment in organelle quality control and metabolic remodeling [25, 26]. This analysis illuminates the key tradeoffs that are governing growth rates in different organisms.

The PLS structure can reveal the growth-promoting and growth-suppressing genes. Independent of the organism, we now ask: what constraints enforce this coordination?

### C. A minimal model predicts mRNA composition

To understand how mRNA composition encodes growth, we define a minimal model of transcription and translation with an empirical constraint (Fig. 4A). Motivated by the tradeoff axes identified by PLS, we classify mRNAs and proteins into three coarse-grained groups: growth-promoting (G^+^), growth-suppressing (G^−^), and other. The G^+^ group increases with growth rate while the G^−^ group decreases with growth rate; notably, the identity of the G^−^ gene set may vary across organisms. In our analysis, the G^+^ group is enriched for ribosomal genes across both organisms, thus we generally identify it with the ribosomal sector moving forward.

**FIG. 4.**
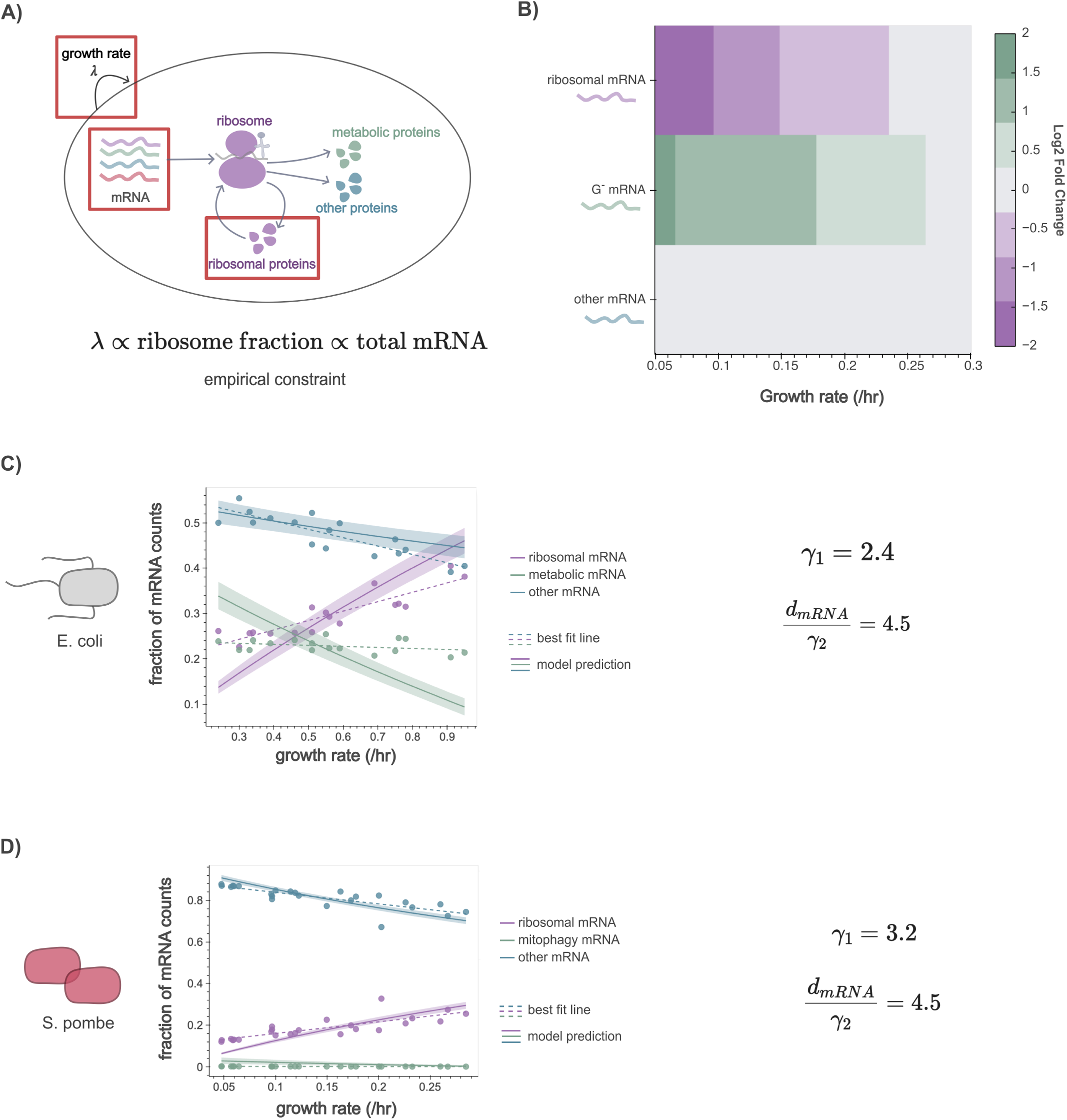
mRNA compositional relationship arises from transcription-translation balance. A) In this model, we focus on the relationship between mRNA composition, translation, and growth. The key constraint is the transcription-translation flux balance constraint that enforces total mRNA and ribosomes together increase with growth. B) Performing hierarchical clustering reveals the existence of a mRNA compositional relationship. Noticeably, the relationship is coordinated with growth. Parameters were chosen to illustrate the qualitative trend; values are provided in the GitHub repository. C) Overlay model predictions with data (and a best-fit line) from *E. coli* (Balakrishnan et al.) and from D) *S. pombe* (Kleijn et al.).

We assume translation rates are constant and distinct for each group, mRNA depletion is dominated by degradation, protein depletion is dominated by growth dilution, and ribosome-mRNA binding follows Michaelis-Menten kinetics. Under these assumptions, the dynamics are:

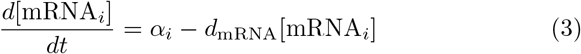

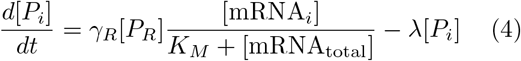

where [mRNA_total_] = ∑_*i*_[mRNA_*i*_] and [*P*_total_] = ∑_*i*_[*P*_*i*_]. Here, *i* indexes the three groups (G^+^, G^−^, or other), *α*_*i*_ is the transcription flux for each mRNA group, *d*_mRNA_ is the mRNA degradation rate, *γ*_*R*_ is the translation rate per ribosome, *K*_*M*_ is the Michaelis-Menten constant, and *λ* is the growth rate. We set [mRNA_other_] to a constant to represent an unchanging portion of the transcriptome.

We incorporate an empirical constraint derived from experimental data – both the ribosomal proteome fraction and total mRNA concentration scale linearly with growth [2,3]:

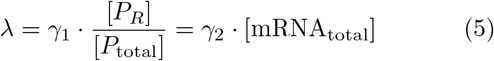

We refer to Eq. (5) as the empirical constraint. While it was originally characterized in *E. coli* and *S. cerevisiae*, we hypothesize these represent universal coordination principles.

By incorporating this scaling law into our minimal framework, the model becomes fully constrained: one relation fixes total transcriptional output as a function of growth rate, the other fixes the ribosomal share of that output, and since *α*_*Q*_is held constant, together they leave no freedom for *α*_*R*_ or *α*_*G*_. Thus, mRNA composition is fully determined by growth rate alone (Supp. Note 1). Consistent with this, the model recapitulates the hierarchical clustering patterns observed experimentally by Gasch et al. (Fig. 4B). Specifically, as growth rate increases, the ribosomal mRNA sector expands while the *G*^−^ mRNA sector contracts.

The identifiable model parameters (*γ*_1_ and *d*_*mRNA*_*/γ*_2_) are biologically interpretable, thus estimating them allows us to test whether the model is quantitatively consistent with experimental measurements and compare translational efficiency across organisms. We used Bayesian modelling and fit the model to transcriptomics data from both *E. coli* and *S. pombe*. The model reproduces experimental trends and magnitudes with some deviations. In *E. coli*, it underestimates the ribosomal mRNA fraction at low growth rates and predicts a steeper decline in metabolic mRNA fraction than observed (Fig. 4C). In *S. pombe*, model predictions closely track the data across all three mRNA groups (Fig. 4D). These *E. coli* discrepancies likely reflect biological details not captured by our minimal assumptions. Despite these limitations, the model successfully predicts the qualitative tradeoff axes. Here, *γ*_1_ is the scaling factor for ribosome allocation, and the ratio *d*_mRNA_*/γ*_2_ is the ratio of the effective mRNA degradation rate to translation rate. In *E. coli, γ*_1_ ≈ 6.4 *×* 10^−4^ s^−1^, whereas in *S. pombe, γ*_1_ ≈ 9.4 *×* 10^−4^ s^−1^. This indicates that, per unit ribosome fraction, ribosomes in *S. pombe* contribute slightly more to growth than ribosomes in *E. coli*. The fitted effective ratio (*d*_mRNA_*/γ*_2_ = 4.5 for both organisms) falls in the expected range derived from experimental values (Fig. 1B). Fitting to proteome data yields similar conclusions (Fig. S4). These results confirm that the specific compositional pattern of ribosomal and metabolic mRNA sectors emerges as a structural requirement of the joint constraint on total mRNA and ribosomal proteome fraction.

### D. Distinct regulatory growth modes emerge from the same model

While the preceding work establishes that mRNA composition is predictive of growth in *S. pombe* and *E. coli*, the regulatory mechanisms underlying this relationship may differ across organisms. Notably, while G^+^ is consistently ribosomal across both organisms, the identity of G^−^ differs — metabolic genes in *E. coli* versus mitophagy-related genes in *S. pombe*. We hypothesize this difference reflects distinct growth-limiting mechanisms. To probe this, we extend our minimal model to explicitly incorporate the molecular function of the metabolic sector (Fig. 5A, S5A).

**FIG. 5.**
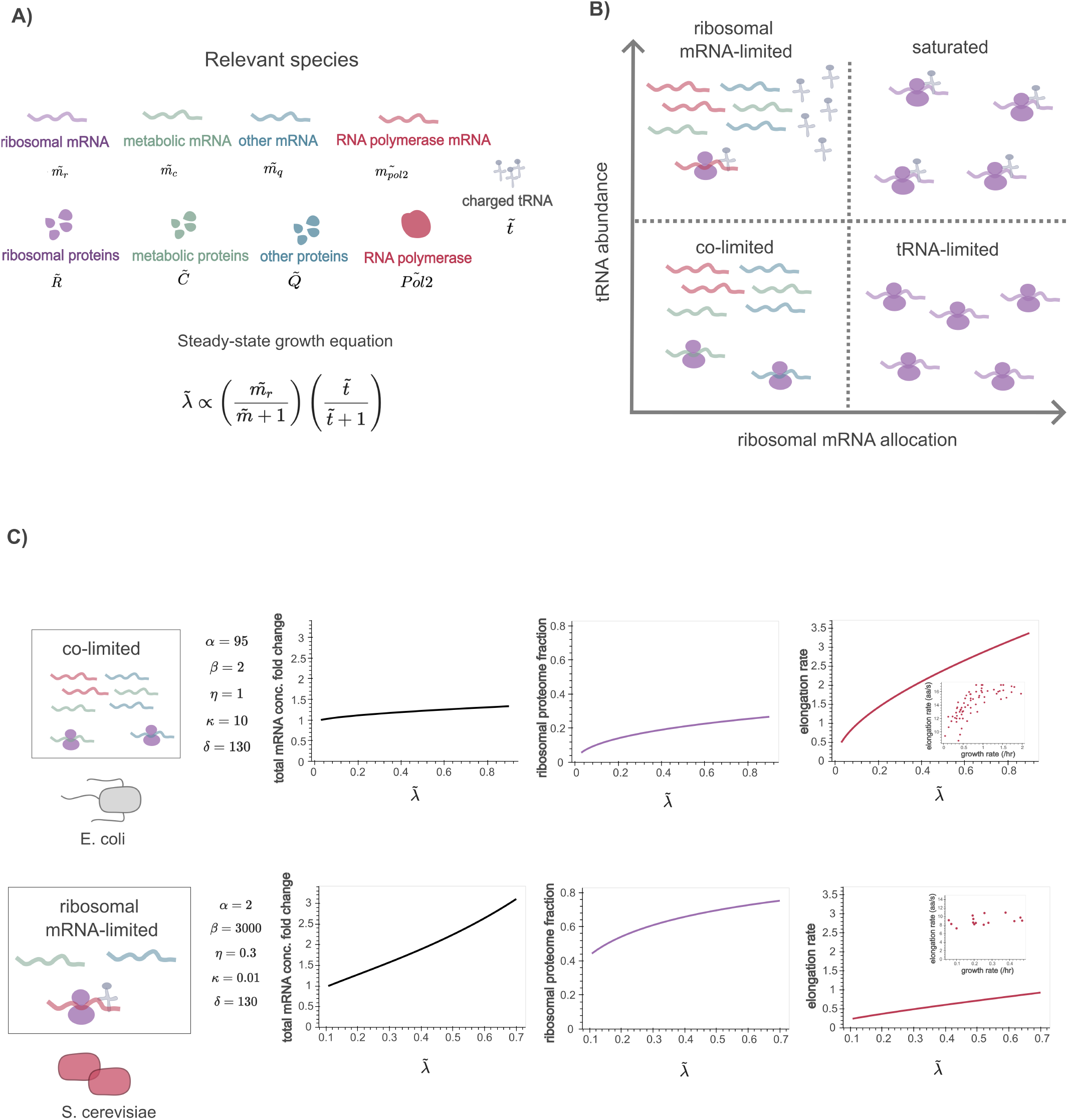
Distinct growth modes emerge from the same model. A) In this model, we consider mRNA species, proteins, and charged tRNA. The steady state growth equation is dependent on the ribosomal mRNA allocation fraction and charged tRNA abundance. B) Four distinct regimes emerge. C) *E. coli* and *S. pombe* display dynamics characteristic of different regimes. Elongation rate plots contain an inset with data (also shown in Fig. 2). Parameters were chosen to qualitatively reproduce empirical trends and lie within ranges motivated by experimental data (from Fig. 1C); the model was not fit to the data.

The previous transcriptomic analysis in this work is based on *S. pombe* and *E. coli*. In addition, we incorporate quantitative measurements from *S. cerevisiae* reported by Gao et al., where scaling relationships between ribosome content, mRNA abundance, and growth rate have been directly characterized [2].

In the extended model, we split mRNA and protein into four categories – metabolic, ribosomal, RNA polymerase (to include details of transcription), and other. We assume transcription rates are distinct for each group and depend on the fraction of polymerases transcribing that group. mRNA depletion is dominated by degradation, protein depletion is dominated by growth dilution, and translation rate is governed by ribosome saturation with two substrates – mRNA and charged tRNAs. The mRNA dynamics follow:

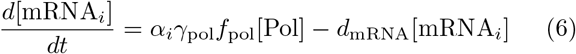

where *α*_*i*_ is the fraction of polymerases transcribing each mRNA sector, *γ*_pol_ is the per-polymerase transcription rate, *f*_pol_ is the fraction of active polymerases, and *d*_mRNA_ is the mRNA degradation rate.

Protein dynamics are described by:

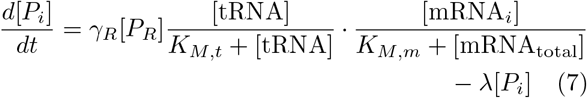

where *K*_*M,t*_ and *K*_*M,m*_ are the Michaelis-Menten constants for charged tRNA and mRNA, respectively, *γ*_*R*_ is the translation rate per ribosome, and *λ* is the growth rate.

We also track charged tRNA:

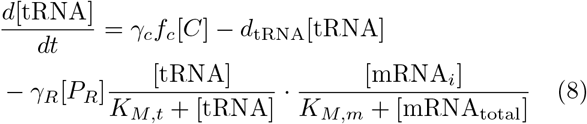

where charged tRNAs are synthesized by catabolic proteins consuming nutrients (*γ*_*c*_ – rate of making charged tRNA from nutrients, *f*_*c*_ – fraction of catabolic proteins making charged tRNAs). Charged tRNAs are depleted by degradation (*d*_tRNA_) and translation. As before, [mRNA_other_] is set to a constant, representing a portion of the transcriptome insensitive to growth.

The elongation rate is defined as a subset of the translation rate:

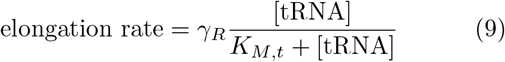

After nondimensionalizing, the steady-state growth equation simplifies to:

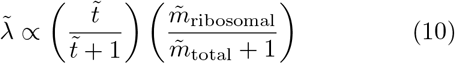

where 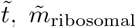, and 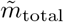 are the nondimensional charged tRNA, ribosomal mRNA, and total mRNA, respectively.

This form reveals four distinct growth regimes (Fig. 5B):

- *Ribosomal mRNA-limited* : low ribosomal mRNA, abundant charged tRNA
- *Co-limited* : both ribosomal mRNA and charged tRNA are low
- *tRNA-limited* : high ribosomal mRNA, low charged tRNA (Fig. S5B)
- *Saturated* : both ribosomal mRNA and charged tRNA are abundant

Comparing the model to experimental data (Fig. 1B) suggests that *E. coli* may reside in the co-limited regime, where both ribosomal mRNA and charged tRNA availability constrain growth. In contrast, *S. cerevisiae* appears in the ribosomal mRNA-limited regime, where tRNAs are abundant but ribosomal mRNA transcripts are limiting (Fig. 5C). This is consistent with observations by Gao et al. showing relatively constant elongation rates across growth rates (Fig. 1D) [2].

By tuning dimensionless parameters inspired by experimentally supported ranges (Supp. Note 2), the model transitions between co-limited and ribosomal mRNA-limited regimes. The parameter *α* = *γ*_*R*_*/d*_mRNA_ represents the protein yield over an mRNA’s lifetime. We find that *α*_*E*. *coli*_ *> α*_*S*. *cerevisiae*_, reflecting a ‘growth-optimized’ prokaryotic strategy characterized by high translation flux. This is supported by experimental data showing elongation rates in eukaryotes are approximately half those of prokaryotes. The lower effective *α* in eukaryotes likely stems from the overhead of nuclear export, splicing, and spatial sequestration, which delays translation and reduces the net protein yield per transcript.

The parameter *β* = *γ*_*c*_*f*_*c*_*/γ*_pol_*f*_pol_ represents the ratio of tRNA synthesis capacity to mRNA synthesis capacity. The finding that *β*_*E*. *coli*_ *< β*_*S*. *cerevisiae*_ suggests that eukaryotes invest heavily in a high supply of charged tRNA. This excessive capacity ensures that tRNA remains saturating and non-limiting, decoupling growth from the specific external condition, thereby making the eukaryotic response intrinsic. In *E. coli*, by contrast, co-limiting tRNA ties growth directly to nutrient-dependent tRNA supply, producing the condition-specific, extrinsic regulation described above.

Finally, the affinity ratio *κ* = *K*_*M,m*_*/K*_*M,t*_ further distinguishes these regulatory modes. The higher *κ*_*E*. *coli*_ indicates that the prokaryotic translation machinery is highly sensitive to the charged tRNA pool, allowing for rapid growth arrest in response to nutrient scarcity. Conversely, the lower *κ*_*S*. *cerevisiae*_ supports a buffered translation system, where growth is less sensitive to tRNA changes.

Altogether, these findings demonstrate that while the underlying biochemical constraints of translation are universal, prokaryotes and eukaryotes occupy distinct regions of the parameter space. The identity of the G^−^ sector may determine which growth regime an organism inhabits. In *E. coli*, where G^−^ corresponds to metabolic genes that directly create charged tRNAs, growth is co-limited by both ribosomal mRNA and tRNA availability. The absence of this coupling in *S. pombe*, where G^−^ reflects mitophagy, may explain why they operate in a ribosomal mRNA-limited regime.

## III. DISCUSSION

This work demonstrates that organism-specific growth tradeoffs can be identified from mRNA composition across environmental conditions in both *E. coli* and *S. pombe*. The axes derived from partial least squares analysis may reflect macro-variables that a cell modulates to navigate a fundamental constraint: the need to partition a finite proteome budget between biosynthesis and stress/catabolism [1,7].

mRNA composition alone is largely sufficient to predict growth rate and determine tradeoffs without explicit knowledge of protein levels or post-translational regulation. This suggests that resource allocation decisions are encoded upstream of protein synthesis, at the level of transcriptional investment across functional sectors. The cell modulates total mRNA abundances and redistributes it across gene sets in a growth-dependent manner.

The empirical observation that total mRNA concentration and ribosomal proteome fractions scale linearly with growth imposes a strong constraint on the existence of a growth tradeoff [1,2]. Our minimal model shows that these constraints necessitate the observed G+/G− structure. Without them, mRNA composition need not encode growth at all. Whether the proportionality is universal and persists across organisms is an open question.

The extended transcription-translation model reveals that *E. coli* and *S. cerevisiae* occupy distinct regulatory regimes despite sharing the same underlying constraints. Prokaryotes appear co-limited by both ribosomal mRNA and charged tRNA availability, while eukaryotes are only ribosomal mRNA-limited. The compartmentalization of eukaryotic cells may allow them to maintain a buffered tRNA pool, channeling growth control through transcription rather than tRNA availability — trading prokaryotic efficiency for regulatory stability.

Several predictions of this framework are falsifiable. First, the framework makes two predictions in any new organism: that a small number of transcriptomic axes should be sufficient to predict growth rate, and that the genes driving those axes should split into opposing sectors. Either poor predictive performance or uninterpretable coefficients would suggest that the linear scaling laws do not hold, that growth is not the dominant selective pressure determining mRNA composition, or the structure of the environments assayed are not sufficiently different or stressful. Second, the regime assignments carry testable consequences. Co-limitation in *E. coli* predicts that perturbations to either charged tRNA or ribosomal transcription should each partially restrict growth, whereas ribosomal mRNA-limited regime of *S. cerevisiae* predicts robustness to tRNA perturbations. Third, introducing an energetic bottleneck by limiting ATP availability may reveal a new growth regime where energy constraints dominate over mRNA and tRNA availability [15,16]. Experiments like these would define the boundaries of the framework and motivate new ones.

Finally, these results can be used to develop a quantitative lens for interpreting dysregulated growth. Cancer and aging cells may alter total mRNA abundance and ribosomal content in ways that violate the linear scaling laws this framework depends on. If so, the identity of genes comprising the growth tradeoff may shift or the relationship between growth rate and mRNA may be entirely different. Mapping pathological cell states onto this framework may reveal whether dysregulation reflects a breakdown of the underlying resource allocation constraints or an escape into otherwise inaccessible growth regimes.

## IV. MATERIALS AND METHODS

All figures and analysis are available at: https://github.com/AnkitaRoychoudhury/StatisticalResourceAllocation

### A. Data acquisition and preprocessing

Transcriptomics, proteomics, and growth rate data for *E. coli* were obtained from Balakrishnan et al. [3]. Table S3 and S4 were used without further processing.

Transcriptomics, proteomics, and growth rate data for *S. pombe* were obtained from Kleijn et al. [11]. Table S1, S3, and S5 were used without further processing.

To assess the relationship between mRNA and protein abundances in *S. pombe*, raw mRNA counts from Kleijn et al. and iBAQ protein abundances from the accompanying proteomics data were matched by gene identifier across shared samples. Genes with zero mRNA counts or zero iBAQ values were excluded. mRNA counts and protein abundances were *log*_10_-transformed, and Pearson correlation was computed in log-log space across all genes and conditions (*R* = 0.61). Only protein-coding genes, as annotated by PomBase, were included in the analysis.

### B. Statistical analysis of mRNA-sequencing and proteomics data

Genes were clustered using agglomerative hierarchical clustering with Ward linkage and Euclidean distance (scipy.cluster.hierarchy). Genes containing missing values across any condition were excluded prior to clustering. A distance threshold of 60 was used to define clusters for *S. pombe* and a distance of 30 for *E. coli*, and genes were ordered by cluster assignment for visualization.

Gene Ontology (GO) enrichment analysis was performed on induced and repressed gene clusters using g:Profiler (Raudvere et al., 2019) via the Python API (gprofiler-official) [24]. Enrichment was tested against GO Biological Process terms using a multiple testing correction method with a significance threshold of FDR *<* 0.05.

PLS regression was implemented using PLSRegression from scikit-learn (v 1.5.1). Prior to fitting, gene expression values were z-scored using StandardScaler. To select the number of PLS components and assess overfitting, data were split into training (75%) and test (25%) sets across multiple random seeds. Training error was estimated via LOO cross-validation on the training set. To establish a null distribution, growth rate labels were permuted 500 times and the full train/test procedure was repeated for each permutation; bootstrap replicates (n=500) were generated by resampling conditions with replacement. Model performance was evaluated using *R*^2^, MSE, RMSE, and MAE across n=1, 2, 3, 4, and 8 components. Two PLS components were selected based on the point at which bootstrapped test error stabilized and remained well-separated from the null distribution.

G^+^ and G^−^ sectors were defined by thresholding the PLS *β* coefficients. For *E. coli*, where PLS was fit separately within each condition class, the coefficient distribution was well-separated and a threshold of zero was used — all genes with positive *β* were assigned to G^+^ and all genes with negative *β* to G^−^. For *S. pombe*, where PLS was fit across all conditions simultaneously, many near-zero coefficients reflect noise rather than biological signal. To identify a principled threshold, a permutation test was performed: growth rate labels were randomly shuffled 1000 times, PLS was refit on each permutation, and the 95th percentile of the resulting null —*β*— distribution was taken as the threshold (*τ* = 6.0810^−5^). Only genes with —*β*— *> τ* were assigned to G^+^ or G^−^; the remainder were excluded from downstream analysis.

### C. Gene set enrichment analysis

Gene set enrichment analysis (GSEA) was performed using the GSEA desktop application (Broad Institute) with genes ranked by *β* coefficient. Gene sets for *S. pombe* and *E. coli* were obtained from the mulea GMT file repository (Ö lbei et al., github.com/ELTEbioinformatics/ GMT_files_for_mulea). Genes were ranked by their *β* coefficient and submitted to the preranked GSEA tool. Terms with FDR q-value *<* 0.25 were considered significant, consistent with standard GSEA guidelines.

### D. Computational modeling

The minimal transcription-translation model partitions the proteome and transcriptome into three functional sectors — ribosomal (R), catabolic/metabolic (C), and other (Q) — and was fit separately to *S. pombe* and *E. coli* data using Bayesian inference implemented in PyMC (v 5.19.1). In all cases, the model was fit simultaneously to observed protein and mRNA sector fractions as a function of growth rate. Posterior sampling used the NUTS sampler with 4 chains, 3000–6000 draws, and 3000–8000 tuning steps. Posterior distributions are summarized as medians with 95% credible intervals. Sector-specific mRNA degradation rates were inferred as free parameters, relaxing the uniform-degradation assumption used in the analytical treatment.

The extended transcription-translation model (Model 2) was analyzed at steady state. Steady-state solutions were obtained numerically using scipy.optimize.root with the hybr method, sweeping across a range of growth rates for parameter sets representative of *E. coli* and *S. cerevisiae*. Parameters were tuned to experimentally supported ranges (see Supplementary Note 2) to identify the distinct growth-limiting regimes described in the main text. All simulations were implemented in Python using NumPy (v 1.26.4) and SciPy (v 1.14.1).

## V. ACKNOWLEDGEMENTS

We thank Arvind Murugan, Elizabeth Jerison, Chris Russo, and the members of the Pincus and Mani labs for their helpful discussions and comments.

We thank the University of Chicago Functional Genomics Facility (RRID: SCR 019196) for bulk RNA sequencing and the University of Chicago Cellular Screening Center (RRID: SCR 017914) for Incucyte instrument access.

This work was supported by National Institutes of Health grant RM1 GM153533 (to D.P. and M.M.); National Science Foundation QLCI QuBBE grant OMA-2121044 (to D.P.); NSF (Grant No. DMS-2235451) and Simons Foundation (Grant No. MPTMPS-00005320) grants to the NSF-Simons National Institute for Theory and Mathematics in Biology (NITMB) (to M.M.); a Chan Zuckerberg Initiative DAF grant (Grant No. DAF2023-329587), an advised fund of the Silicon Valley Community Foundation (to M.M.); and NSF (Grant No. PHY-1748958) and Gordon and Betty Moore Foundation (Grant No. 2919.02) grants to the Kavli Institute for Theoretical Physics (to M.M.).

## VI. SUPPLEMENTARY NOTES

### A. Supplementary Note 1: mRNA composition is determined by the constraints in the minimal model

In the minimal transcription-translation model, steady-state analysis reveals:

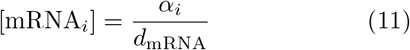

where *i* represents the sectors – G^+^ (ribosomal, represented by *R*), G^−^, and other (represented by *Q*). Then:

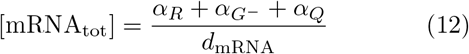

The *d*[*P*_*R*_]*/dt* equation at steady state provides the growth equation:

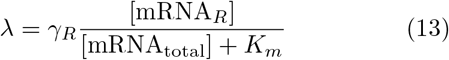

Equating with the other steady-state protein equations, we find that mRNA and protein fractions are linked such that:

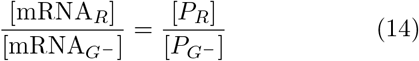

and

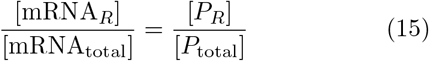

Thus, each protein fraction mirrors the corresponding mRNA fraction.

Adding the constraint *λ* = *γ*_2_[mRNA_total_], we can write:

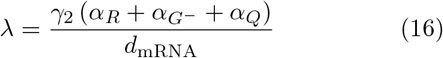

or equivalently:

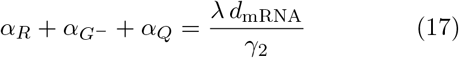

This indicates that the sum of transcription fluxes for all mRNA species is capped at a value set by the growth rate and two constants.

Adding the constraint *λ* = *γ*_1_[*P*_*R*_]*/*[*P*_total_], we can write:

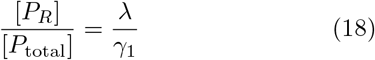

Thus, the fraction of ribosomal proteins is set by the growth rate. Recall that since

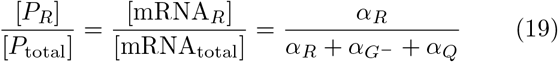

this constrains what *α*_*R*_ must be. Since *α*_*Q*_ is set to a constant, *α*_*R*_ is determined by the second constraint, and the sum 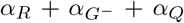 is set by the first constraint, each mRNA sector’s transcription flux is fully determined. Thus, mRNA composition is determined by the joint empirical constraint.

### B. Supplementary Note 2: Nondimensionalization of extended transcription-translation model

Define *τ* = *t* · *d*_mRNA_, 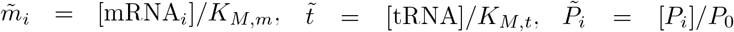, where *P*_0_ = *K*_*M,m*_ *d*_mRNA_*/γ*_Pol2_*f*_Pol2_.

The nondimensional parameters are defined as:

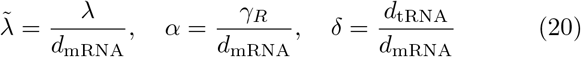

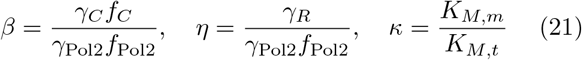

The full set of dimensionless equations becomes:

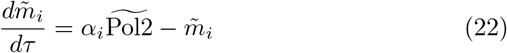

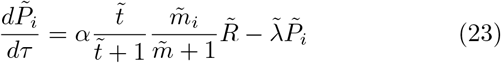

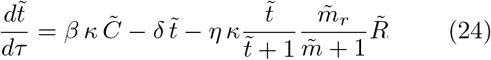

where 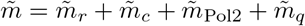.

The nondimensional variables represent key cellular balances. We derive ranges for these parameters from experimentally measured quantities.

Time is normalized by the mRNA degradation rate, so *τ* represents time in units of mRNA lifetime. Similarly, growth 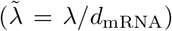 is expressed in units of mRNA lifetime.

*α* = *γ*_*R*_*/d*_mRNA_ is a balance between translation and mRNA degradation rate – it can be interpreted as the amount of protein produced per mRNA lifetime. When *α >* 1, translation is fast relative to mRNA turnover, which is the regime we expect cells to occupy. Using evidence from bacteria that translation rates range from 0.03–0.06 s^−1^ and mRNA degradation rates range from 0.046–0.69 min^−1^, we estimate *α*_*E*. *coli*_ ≈ 3–78. Data from yeast indicates translation rates of 0.01–0.03 s^−1^ and mRNA degradation rates of 0.011–0.069 min^−1^, giving *α*_*S. cerevisiae*_ ≈ 9–164.

*β* = *γ*_*C*_*f*_*C*_*/γ*_Pol2_*f*_Pol2_ is a balance between charged tRNA and mRNA production. When *β* is high, there is more charged tRNA relative to mRNA. This parameter titrates the system between a tRNA-limited regime and a ribosomal mRNA-limited regime. While estimates of *γ*_*C*_, *f*_*C*_, and *f*_Pol2_ are scarce, typical transcription rates can be used to estimate *γ*_Pol2_. Bacteria have transcription rates of 0.01–0.1 s^−1^, while yeast transcription rates have been measured at ~ 0.04 s^−1^.

*η* = *γ*_*R*_*/γ*_Pol2_*f*_Pol2_ is the ratio of the translation rate per ribosome to the rate of mRNA synthesis per Pol2. This ratio is balanced when mRNAs are consumed by translation as fast as they are produced. *η <* 1 indicates excess mRNA relative to ribosome translation capacity. Assuming a constant *f*_Pol2_, translation and transcription rates in bacteria and yeast can be used to estimate *η*.

*κ* = *K*_*M,m*_*/K*_*M,t*_ represents the ratio of the mRNA and tRNA saturation scales. This value can push the system between mRNA-limited and tRNA-limited regimes. Lastly, *δ* = *d*_tRNA_*/d*_mRNA_ is the ratio of tRNA stability relative to mRNA stability. *δ* ≫ 1 represents a regime where charged tRNA is significantly more stable than mRNA.

---

**FIG. S1.**
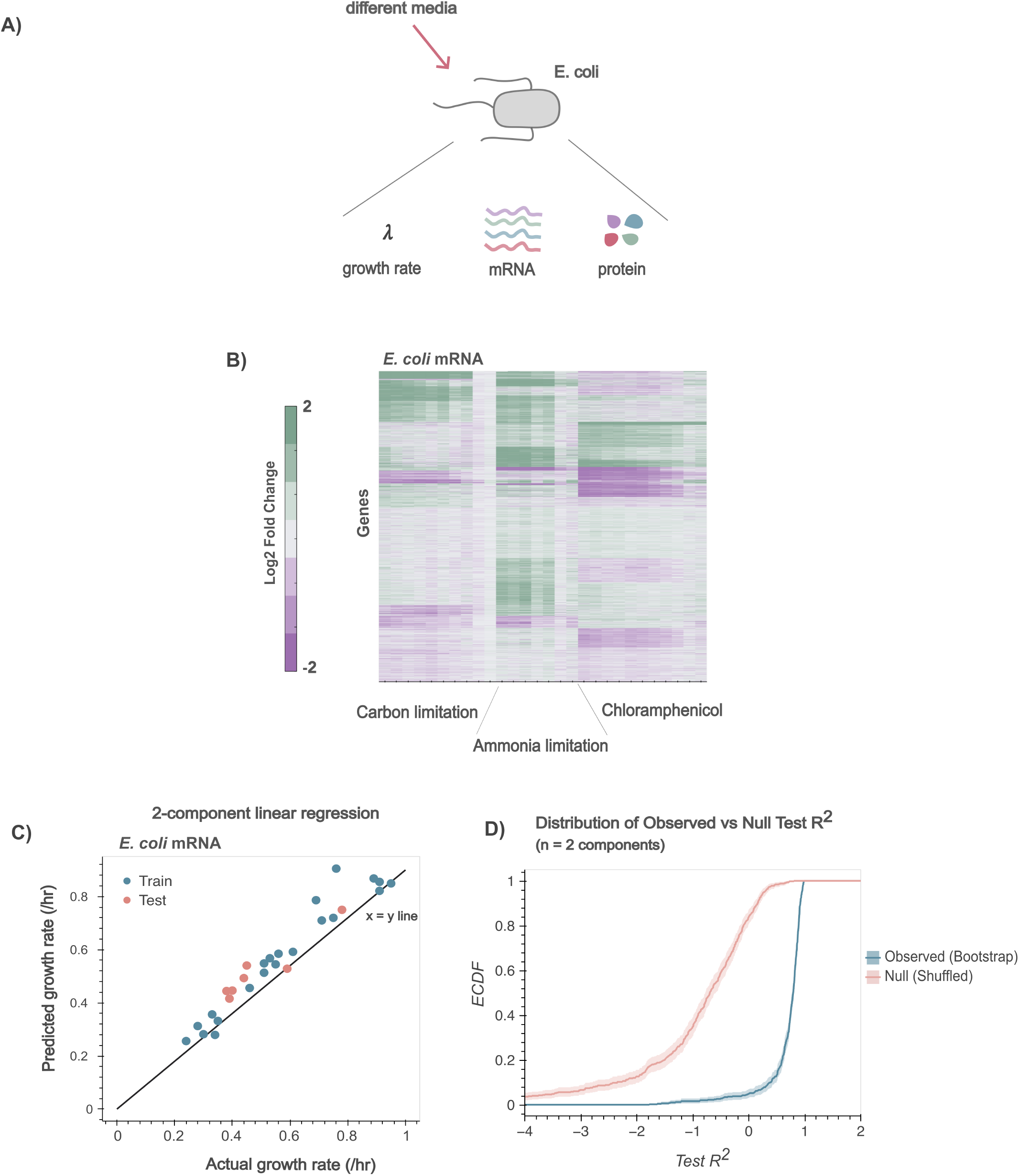
Data from *E. coli* (Balakrishnan et al.) shows the relationship between mRNA composition and growth. A) Balakrishnan et al. collected data in *E. coli* grown in media with different limitations - carbon, ammonia, and ribosome. Proteomics, mRNA sequencing and growth data were collected. B) Hierarchical clustering performed on mRNA counts. C) PLS on the mRNA-sequencing data shows that growth information is indeed encoded in the transcriptome. D) Model validation via permutation testing. ECDF of test *R*^2^ values for the *n* = 1 component model across 500 bootstrap replicates. The observed bootstrap distribution is significantly shifted from the null distribution of shuffled labels, confirming the model captures genuine biological signal. 95% confidence intervals are shaded. X-axis is truncated to [−4, 2] for visual clarity.

**FIG. S2.**
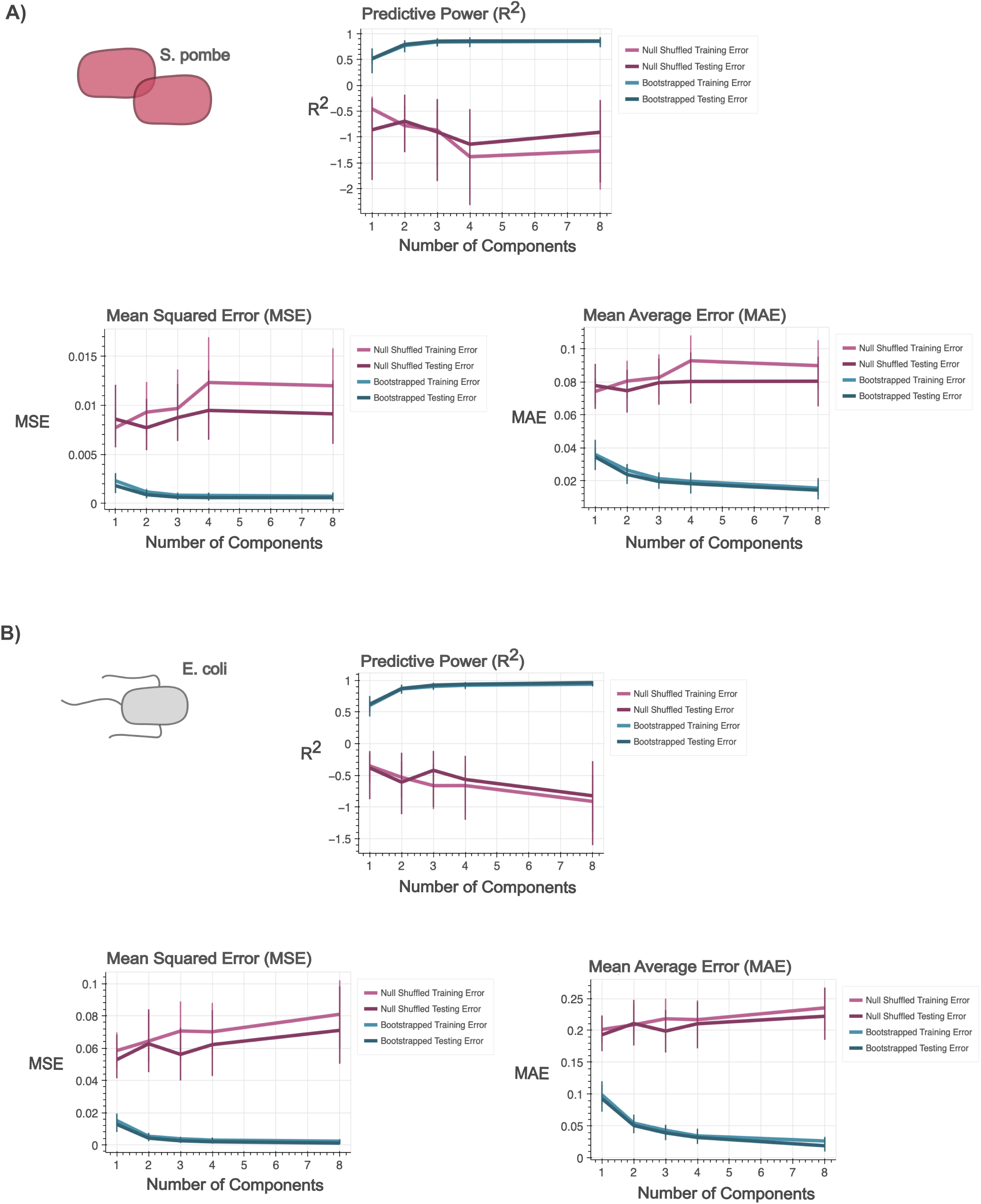
Selection of PLS components via cross-validation. Model error (MSE, RMSE, MAE) and predictive power (R2) as a function of the number of PLS components. The observed model (blue) demonstrates a reduction in error at n=2 components, with stable performance across additional components. In contrast, the null model (maroon), generated via label shuffling, maintains high error and poor predictive power regardless of the number of components used. Error bars represent the standard deviation across 500 bootstrap replicates and cross-validation splits. A) Data from Kleijn et al. measured in *S. pombe*. B) Data from Balakrishnan et al. measured in *E. coli*.

**FIG. S3.**
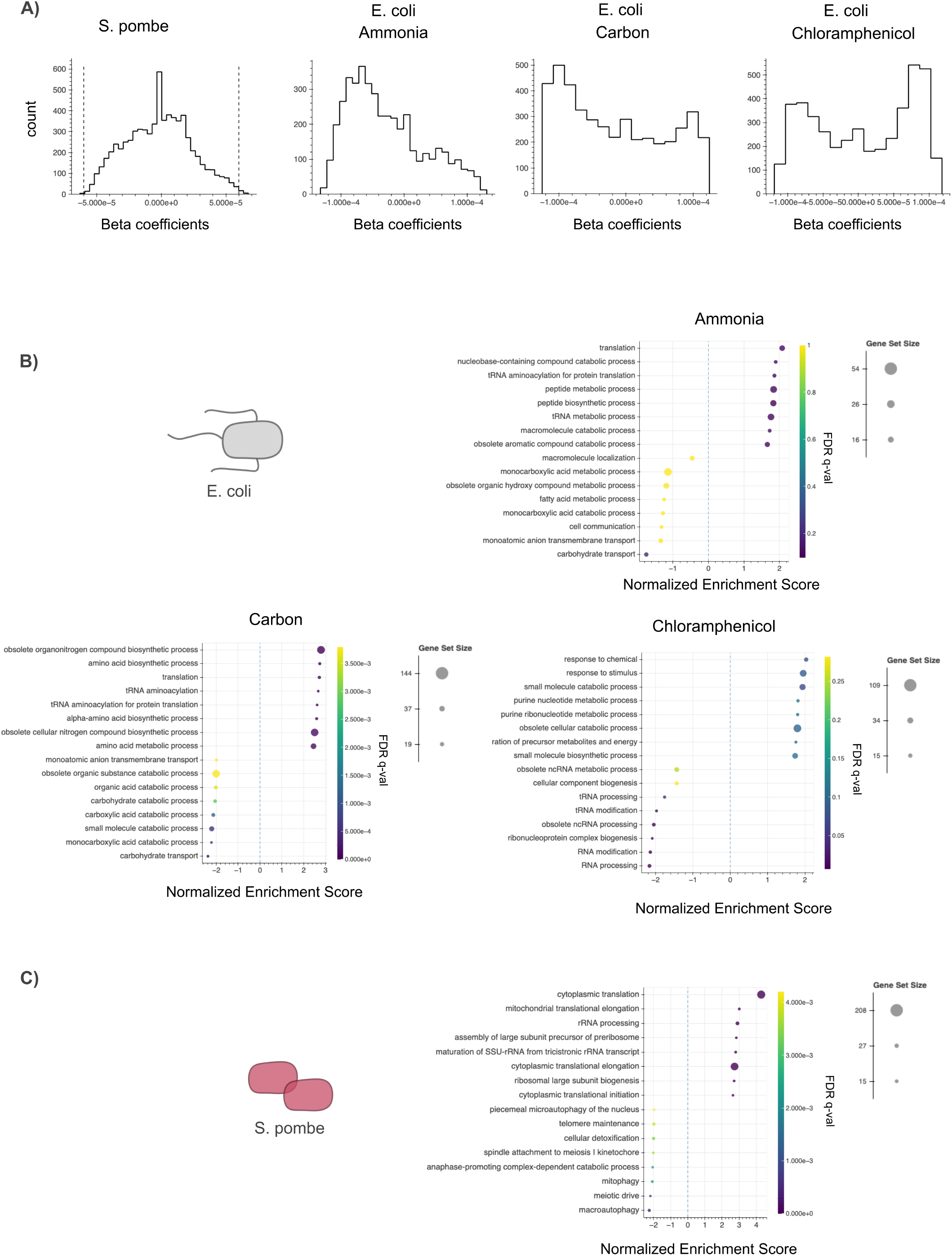
Beta distributions and details of the GSEA. A) Histograms of beta distributions. *E. coli* distributions have the threshold (*τ* = 6 ∗ 10^−5^) represented by a gray dotted line. B) Top GSEA results for each *E. coli* conditions. C) Top GSEA results for the entire *S. pombe* dataset.

**FIG. S4.**
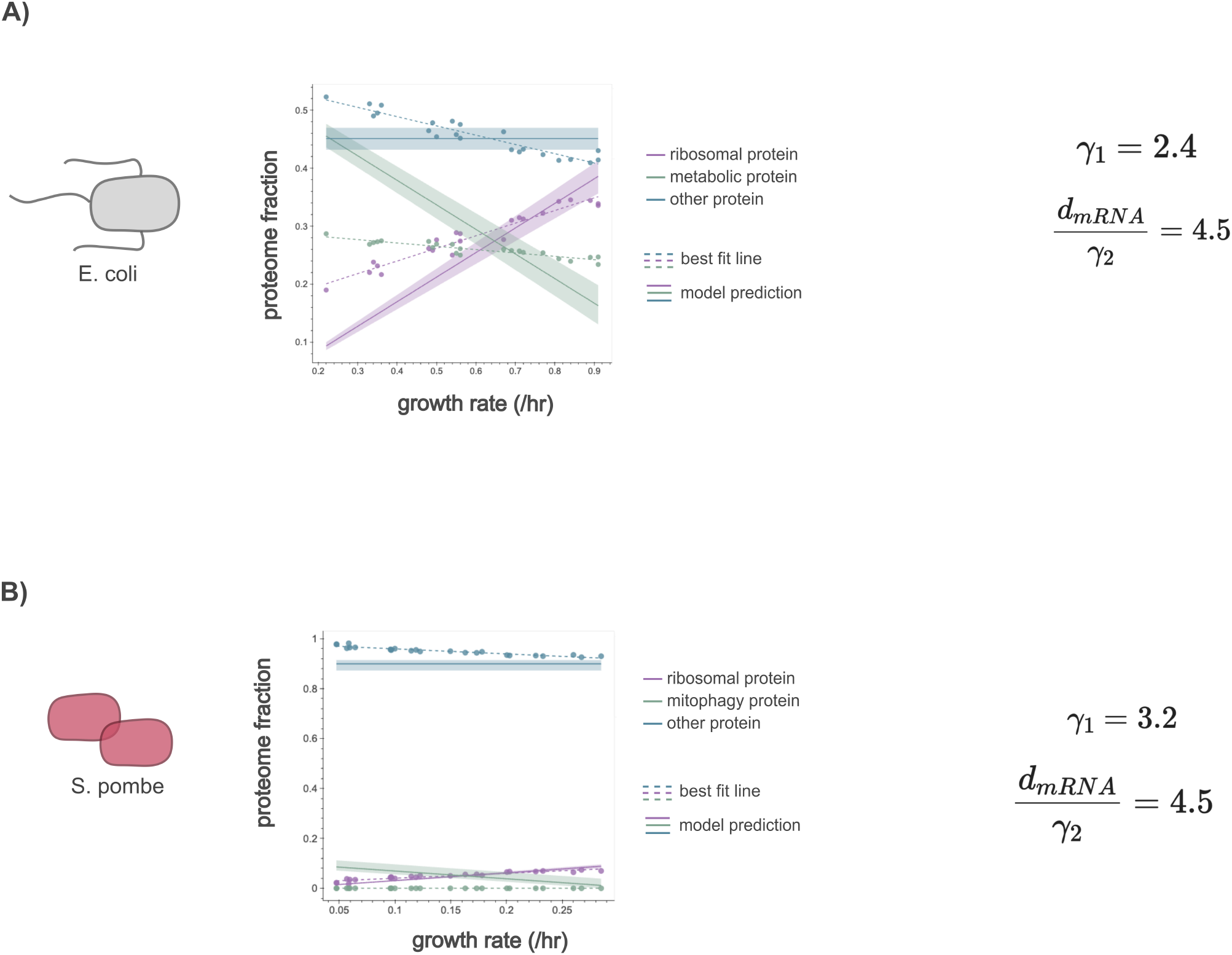
Fitting protein data over minimal transcription-translation model. A) Overlay model predictions with proteomics data (and a best-fit line) from *E. coli* (Balakrishnan et al.) and B) *S. pombe* (Kleijn et al.).

**FIG. S5.**
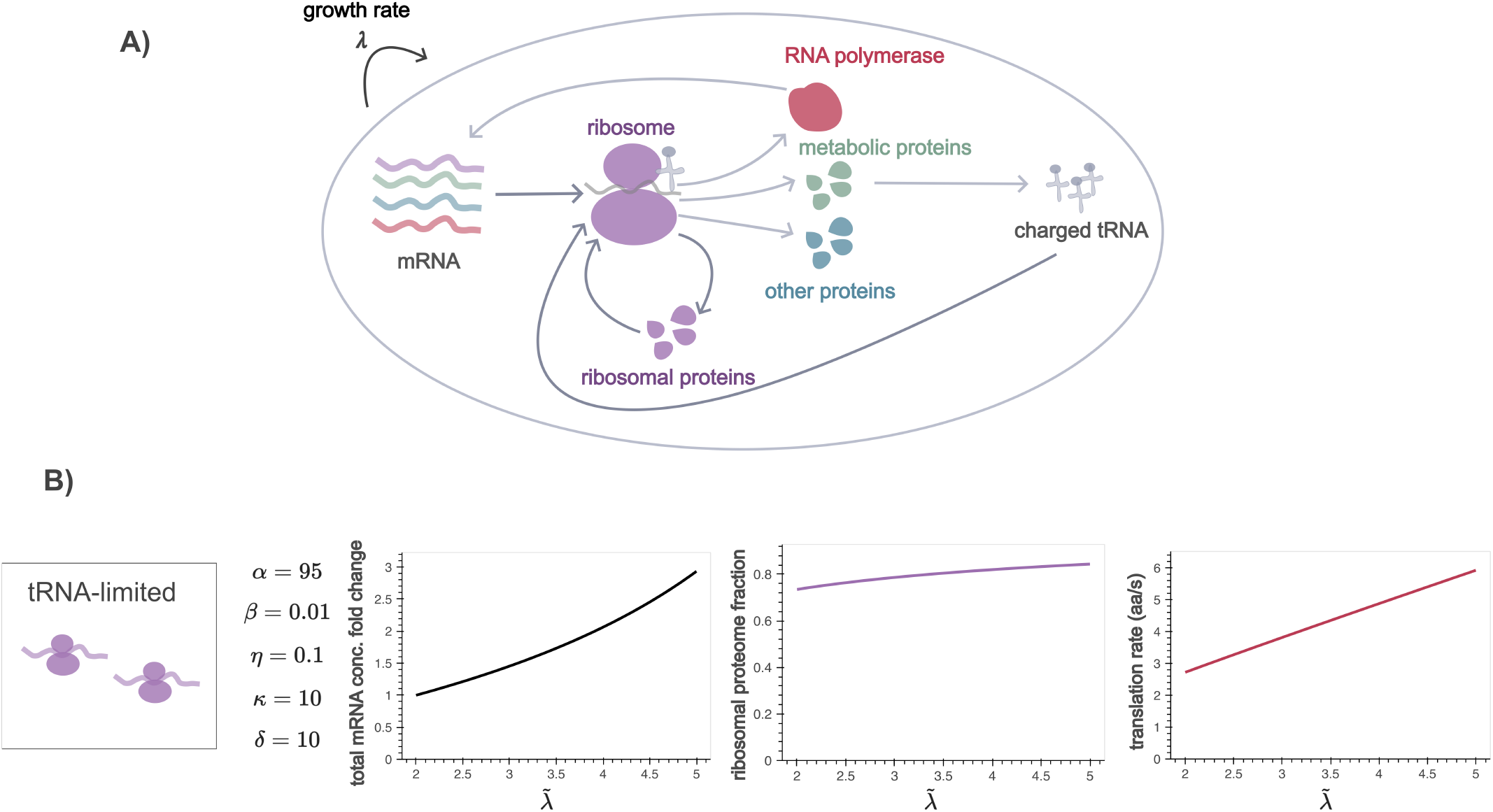
Exploring the tRNA-limited regime in the extended transcription-translation model. A) An example set of parameters displaying the dynamics for the tRNA-limited regime. Parameters were chosen to lie within ranges motivated by experimental data.

